# Phantom- and simulation-based validation of combined diffusion relaxometry in ex vivo ADRD white matter

**DOI:** 10.64898/2026.04.27.721146

**Authors:** Andrea Jacobson, Amaya Murguia, Scott D. Swanson, Jon-Fredrik Nielsen, Jeffrey A. Fessler, Navid Seraji-Bozorgzad

## Abstract

**Purpose:** In principle, combined T_2_–Diffusion (D) MRI has the microstructural and chemical sensitivity to detect axonal and myelin water changes in Alzheimer’s disease and related dementias (ADRD), but its practical implementation may be hindered by demanding hardware requirements. This work assesses the feasibility and accuracy of T_2_–D for ex vivo analysis of WM lesions in ADRD tissue.

**Methods:** A thawed ex vivo brain sample from the Michigan Brain Bank and a T_2_–D phantom were scanned at 7T using a combined diffusion relaxometry (CDR) sequence. A non-negative least squares (NNLS) conventional data processing pipeline was used to disentangle water pools with unique T_2_–D signatures. Simulations examined the effects of minimum TE and SNR on recovery of myelin water (short T_2_, slow diffusion).

**Results:** Across tissue types, T_2_–D data consistently resolved three spectral components. Phantom experiments showed detection of short T_2_ and slow diffusion features similar to those observed in ADRD ex vivo tissue, and confirmed CDR’s ability to accurately resolve multiple components. Simulations indicated reliable T_2_–D recovery for myelin with SNR *>* 30 dB and minimum TE *<* 25 ms.

**Conclusion:** Strong T_2_ and D weighting could be combined to capture the expected axonal, myelin, and extracellular (EC) regions in T_2_–D space. The observed short-T_2_, restricted-D components are therefore unlikely to be artifacts and instead support interpretations as physically meaningful myelin and axonal water signatures.

## 1 INTRODUCTION

ADRDs are leading causes of morbidity in aging populations, ^1^ characterized by cognitive decline and functional impairment. ^2,3^ While neuronal loss in gray matter (GM) is a hallmark of Alzheimer’s disease (AD), the contribution of white matter (WM) lesions has been increasingly recognized. ^4,5,6,7^

Clinically, WM hyperintensities on T_2_ FLAIR imaging are used as sensitive markers of WM lesions, ^8,9,10^ and diffusion weighted imaging is widely used to detect microstructural abnormalities. ^11,12^ However, these techniques individually cannot readily separate axonal, extracellular, and myelin water pools. ^13,14,15^

Combined diffusion relaxometry (CDR) uses both T_2_ and diffusion measurements to distinguish between biological water environments. ^16^ CDR data are commonly represented as a two-dimensional T_2_–D spectrum, obtained via an inverse 2D Laplace transform (ILT),which decomposes the measured signal into multiple T_2_–D pairs (or spectral components). ^17,18^ They can represent the distinct signal contributions associated with different microstructural and chemical environments in tissue. This technique has found applications ranging from obstetrics to neurology. ^19,20,21,22^

T_2_–D imaging is conceptually well suited for separating WM water pools of interest in AD, but the components of greatest interest impose competing acquisition requirements. Sensitivity to axonal water benefits from strong diffusion weighting (high *b* values), whereas detecting myelin water requires short TEs. However, achieving high *b* values often requires longer diffusion-encoding gradients, thereby increasing the minimum achievable echo time (TE_min_). ^23^ These inherent tradeoffs are further complicated by the achievable signal-to-noise ratio (SNR), as noise can corrupt the ability to resolve restricted diffusion coefficients. ^24^ These considerations raise two practical questions: 1) should our T_2_–D fitting routine exclude the myelin water peak, which could affect the resolved axonal and EC component T_2_–D values and 2) what TE_min_ and SNR are required to reliably capture and disentangle both myelin and axonal water spectral components?

In this work, we analyzed CDR data from an ADRD ex vivo brain sample and validated our observed spectral components using two complementary tools: (1) a phantom designed to systematically mimic components of interest, and (2) simulations of CDR datasets with varied acquisition parameters (TE_min_) and hardware constraints (SNR).

## 2 METHODS

### 2.1 Ex Vivo Tissue Selection and Scanning

Our process and procedures were compliant with and conducted with the approval of our university’s Institutional Biosafety Committee (IBC). A frozen ex vivo post-mortem brain from a donor who had a pathologically confirmed diagnosis of AD and cerebral atherosclerosis was selected from the Michigan Alzheimer’s Disease Research Center (MADRC). Frozen tissue was used to avoid formalin crosslinking, which is known to affect T_2_ and D. ^25,26^ A sample containing both WM lesions and normal appearing white matter (NAWM) was dissected, guided by the pre-mortem T_2_ FLAIR scan, and prepared as described by Murguia et al.. ^27^

The sample was placed in a 40 mm Millipede quadrature coil and inserted into a 7.0 Tesla NMR/MRI small animal scanner (Varian/Agilent, Walnut Creek, CA, USA) with 40 mT/m gradients with a 115-mm inner diameter. The temperature was approximately 15−20^°^C to preserve the sample while scanning. Two rounds of voxel-wise, 3D gradient-echo shimming were performed.

A two-shot center-out spin-echo EPI scan was performed using 380 pairs of *b* values (19 values, from 1.49 to 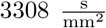) and TEs (20 values, from 18.92 to 189.2 ms). Diffusion weighting was strengthened by using all three directional gradients simultaneously, achieving a gradient amplitude of 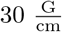, 4 ms duration, and 12 ms separation. The in-plane resolution was 0.547×0.547 mm (FOV = 35 × 35 mm, matrix size = 64 × 64). The TR was 2000 ms, and the scan time was 50 minutes.

### 2.2 Phantom Development and Scanning

A phantom was designed to emulate multi-component systems (see Gatidis et al.). ^28,29^ The phantom contained 15, 5 mm NMR tubes with different concentrations of polyethylene glycol 3350 (PEG) and gadolinium (Gd) contrast agent gadobenate dimeglumine MultiHance (Bracco Diagnostics Inc., Princeton, NJ, USA). The concentration of Gd changed the T_2_ values (5-100 ms) and the addition of 100 mM of PEG resulted in two different D values (approximately 1.5 and 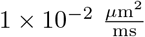 (Figure 1 A).

**FIGURE 1.**
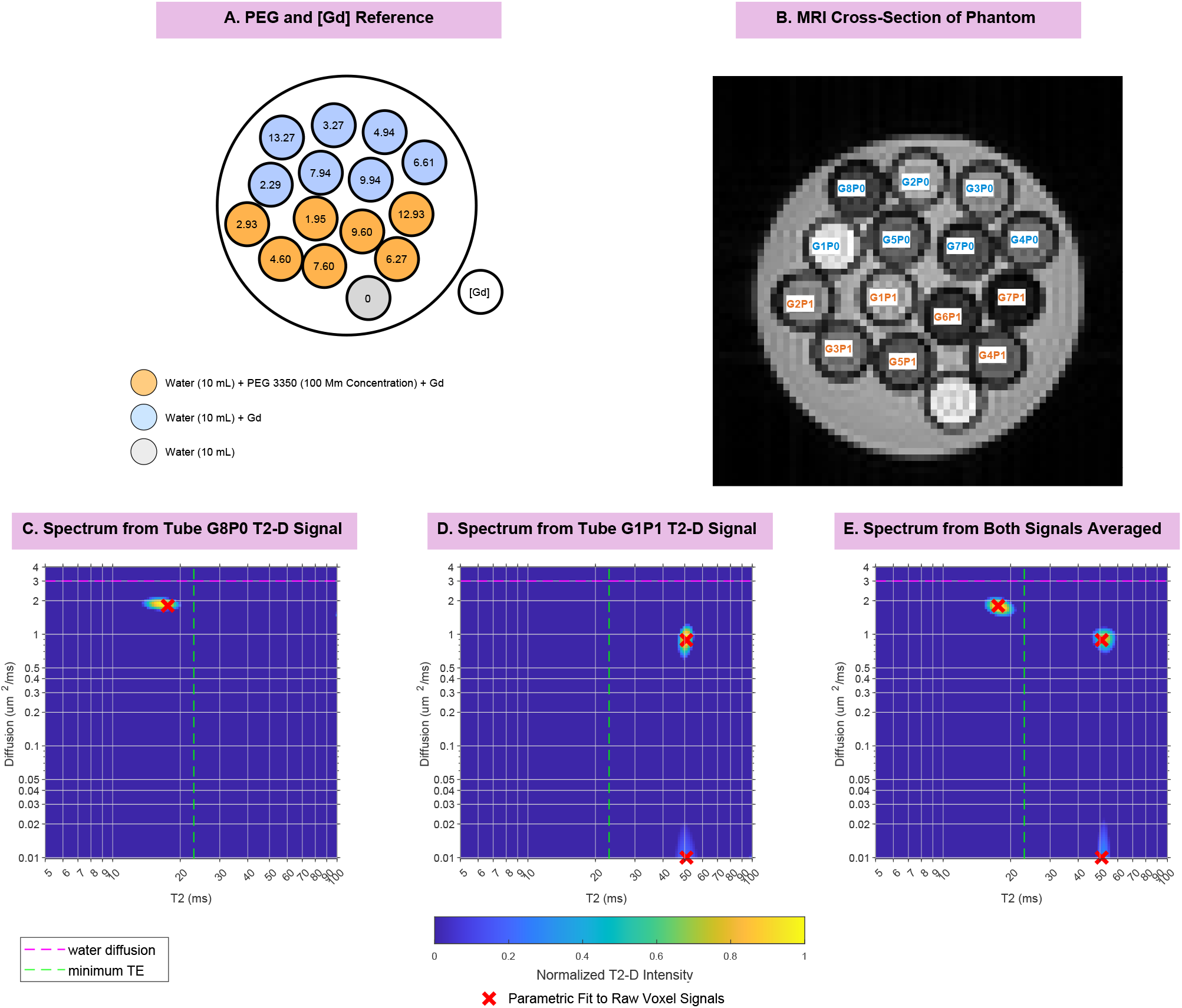
Physical phantom for generating ground truth T_2_–D spectra and synthetic multi-component voxels. A) Phantom Composition: PEG+Gd phantom comprising 15 tubes with known and varying diffusion (D) and T_2_ values. Gd concentration ([Gd]) is labeled at the center of each tube; orange shading indicates the presence of 100 mM PEG 3380. B) MRI profile: T_2_-weighted MR image of the phantom demonstrating signal variation with increasing [Gd]. Tubes are labeled G1–G8 in order of increasing [Gd]; the suffix P1 or P0 denotes the presence or absence of PEG, respectively. C-D) Example D–T_2_ spectra. The individual signal’s mono-or bi-exponential parametric fits are marked with 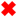’s. E) Composite spectrum construction: Spectrum of “synthetic” voxel, created by adding the ROI-averaged MRI signals from vials in (C) and (D). The three 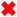 marks are translated from C and D for reference. This incremental combination enables systematic construction of tissue-like spectral components with ground truth T_2_–D values.

A spin echo (non-EPI) T_2_–D sequence was performed using 132 pairs of *b* values (11, from 0 to 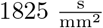) and TEs (12, from 23 to 133 ms) (Figure 1 B). The diffusion weighting used a maximum gradient amplitude of 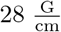, duration of 6 ms, and separation of 11.3 ms. The in-plane resolution was 0.547 × 0.547 mm. The TR was 6000 ms, and the scan time was approximately 14 hours.

### 2.3 Data Processing

#### Non-negative least squares T_2_–D fitting

NNLS fitting ^17,18^ was applied to all datasets. Regions of interest (ROIs) were manually selected in MATLAB from the T_2_–D MR images. To avoid bias from the spatially varying phase in complex signal averaging, phase correction was applied by computing the circular mean phase over the ROI voxels and multiplying each complex image by the corresponding conjugate phasor to rotate the mean ROI phase toward zero. The real components of the corrected signal intensities were then averaged within each ROI and stored as the measured signal vector, ***y***.

The 2D spectrum ***x*** was estimated by solving

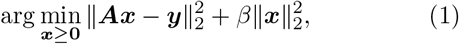

where ***A*** is a dictionary containing the signal evolutions. Each column of ***A*** corresponds to a unique (T_2_, D) pair, and each row corresponds to a specific (TE, *b*) acquisition, with entries given by ^17^

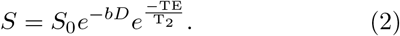

The T_2_ and D ranges used to construct ***A*** were informed by prior observations and previous studies. ^21,27,30^ A total of 120 T_2_ values and 121 diffusion coefficients were included.

Non-negativity constraints and Tikhonov regularization were imposed to restrict solutions to physically meaningful values and to improve robustness to noise and numerical instability. ^31,32^ The regularization parameter *β* was selected in the range 10^*−*6^–10^*−*5^, guided initially by cross-validation and refined based on consistency with prior NMR experiments and the ability to resolve distinct components. Solutions were obtained using a built-in NNLS solver (MATLAB R2023a (MathWorks, Natick, MA, USA).

#### Ground truth D and T_2_: 1D parametric fitting

As the 2D NNLS fitting routines are ill-posed by nature, we needed to calculate ground truth measures for use in the phantom experiments. The one-and two-component phantom tubes were analyzed using mono or bi-exponential curve fitting, respectively. We used all acquired TEs and the first *b* value to estimate T_2_, and all acquired *b* values and the first TE to estimate D. The resulting coefficients, or parametric component fits, were treated as ground truth values.

### 2.4 Phantom Applications for WM Spectra Validation

Components in the 2D spectrum characteristic of both lesioned and normal-appearing white matter (NAWM) tissue were emulated with the phantom by combining MR signals from individual tubes with known diffusion (D) and T_2_ coefficients. Tube signals were averaged together incrementally, allowing the contribution of each spectral feature to be examined in isolation.

#### Example phantom application

Here, we describe how the phantom can be used. In this example, we can set our goal arbitrarily to construct a spectrum that will include components with the following T_2_–D features:

- T_2_ shorter than the TE_min_
- highly restricted D with a long T_2_
- less restricted D with a long T_2_

First, reference spectra and ground truth parametric fits were calculated individually (section 2.3). Here, tube G8P0 was used to get a short T_2_ component (Figure 1C), and tube G7P1 was used for the other two components (Figure 1D). Next, the ROI-averaged signals from both tubes were combined and analyzed to produce a composite spectrum (Figure 1E). Finally, all three spectra (two individual fits and composite) can be visually compared to assess any spectral artifacts and to confirm the component placement is accurate and consistent between the spectra.

#### NAWM and lesion

To emulate NAWM and lesion spectral features, two different pairs of tubes were used. Tube selection was guided by representative NAWM and lesion spectra. Tubes G3P1 and G7P1 were used for NAWM, and tubes G2P0 and G5P1 for lesions.

### 2.5 Simulations for Investigating the Effects of Experimental Noise and Minimum TE

Data for the T_2_–D experiment was simulated for a myelin water pool (T_2_ = 7 ms and 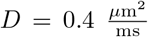). The same number of TEs, the maximum TE (209 ms), and the *b* values from the ex vivo tissue scans were retained for all datasets, but each simulation used a different combination of TE_min_ and SNR. The TE_min_ values ranged from 5 to 50 ms, and SNR values from 10 to 50 dB. A total of 120 T_2_–D spectra were generated using the processing pipeline described in section 2.3.

To assess both the precision and accuracy of the resolved spectral component locations, we defined metrics called “spread” and “bias,” respectively. Before calculating these metrics, our spectral peaks were thresholded to exclude points with a value (spectrum intensity) below 50 percent of the maximum intensity.

#### Bias

The centroid of the myelin water spectral compo-nent was calculated as

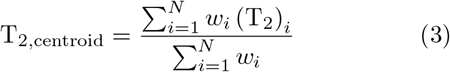

along the T_2_ axis, and likewise along the D axis. Here, *w*_*i*_ is the weight at pixel *i*, T_2,*i*_ is the T_2_ coefficient at pixel *i*, and *N* denotes the number of pixels within the range where the peak is expected to appear. These centroids were subtracted from the true coefficients to obtain a bias metric. For example, in the T_2_ dimension,

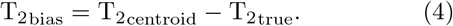

#### Spread

The spread, or width, of the spectral peaks in the D and T_2_ dimensions was determined by first identifying the longest connected sequence of spectral bin in each direction. The spread was then calculated as the difference between the maximum and minimum D and T_2_ values. The minimum spread that could be assigned was the spectral bin width. The data are displayed as spread and bias maps using contour plots.

## 3. RESULTS

### 3.1 Ex vivo tissue

The spectra derived for NAWM, lesions, and GM consistently showed three peaks (Figure 2). We will refer to the tissue spectral features as Components 1-3, defined by their T_2_–D properties as such:

- Component 1: Short T_2_, appearing below the TE_min_, speculated to be related to myelin water
- Component 2: Longer T_2_, speculated to be extracellular (EC) water
- Component 3: Same T_2_ as Component 2, but with a restricted diffusion coefficient located near the lower boundary of the signal evolution dictionary, speculated to be axonal water.

**FIGURE 2.**
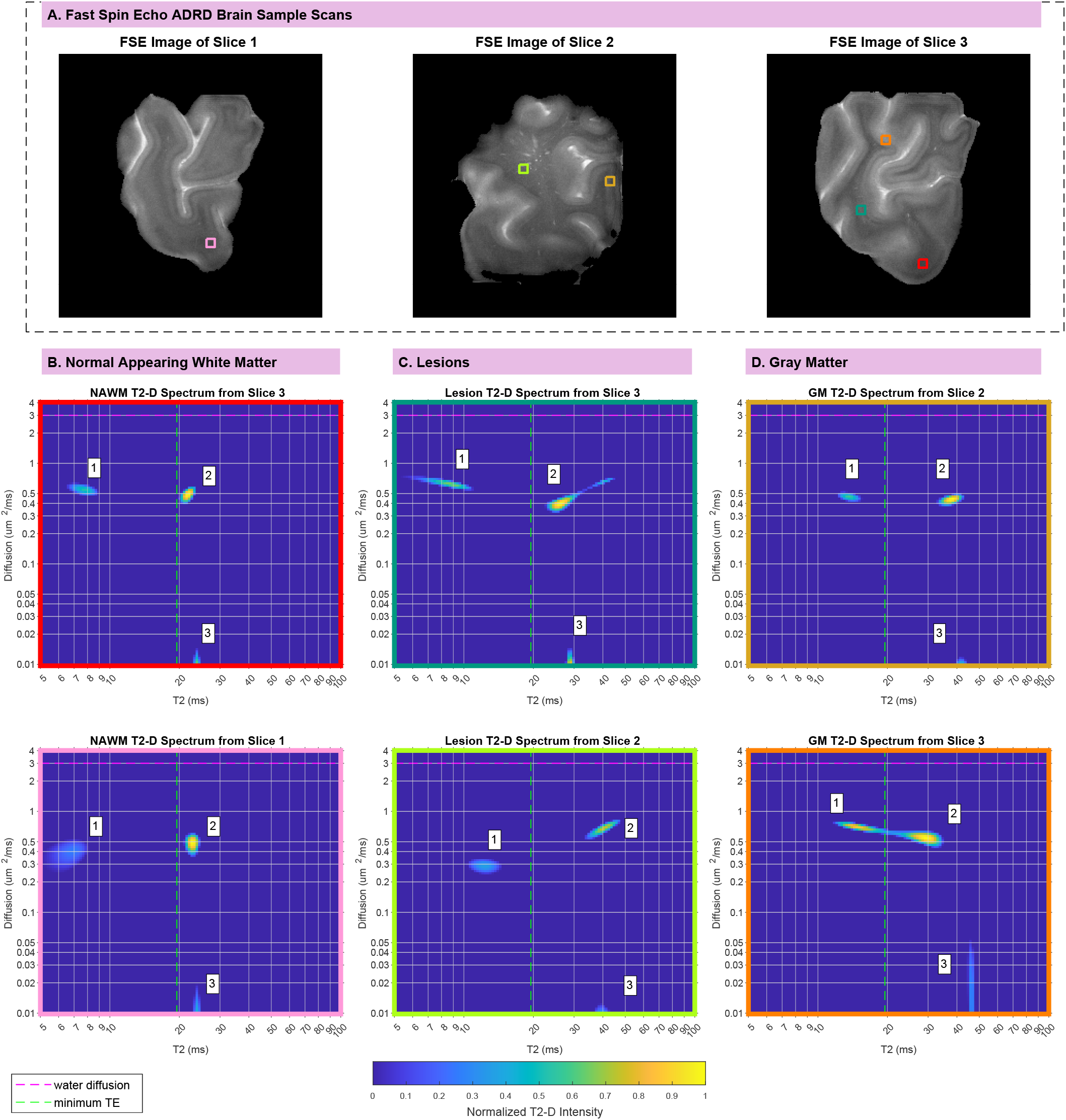
Observed T_2_–D spectral components in a brain tissue sample from an ADRD patient donor. A) Fast spin echo images of different slices of the brain tissue sample. The border color of each ROI matches that of its corresponding T_2_–D spectrum. T_2_–D spectra for B) normal appearing white matter (NAWM), C) lesions, and D) gray matter (GM). Across all spectra, three consistent peaks are observed: (1) component with T_2_ below the TE_min_, (2) component with T_2_ above the TE_min_ and closer to free diffusion, and (3) component with a highly restricted diffusion coefficient. These observed spectral components could describe biological features.

#### Normal Appearing White Matter

Components 2 and 3 had T_2_s between 25 - 30 ms (Figure 2 B). Component 1 had a T_2_ value around 6 - 10 ms, less than half the TE_min_. Components 1 and 2 had diffusion coefficients between 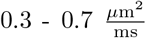. The fitting routine pinned component 3 consistently at the slowest diffusion coefficient,.

#### Lesions

Components 2 and 3 had T_2_s between 25 - 50 ms (Figure 2 C). Component 1 appeared closer to the TE_min_ than was seen with the NAWM. The diffusion coefficients for all three peaks were similar to the NAWM results.

#### Gray Matter

The spectra for gray matter (GM) were similar to those of the lesions; however, on average, all three peaks appeared at slightly longer T_2_s.

### 3.2 Phantom

#### Lesion spectrum validation

We were able to use phantom tube G2P0 (Figure 3B) to derive a component with a T_2_ similar to that of the lesion’s Component 2 (Figure 3A). Additionally, we were able to replicate the diffusion coefficients, which varied by approximately two orders of magnitude, for the lesion’s Components 1 and 3 with tube G5P1 (Figure 3C).

**FIGURE 3.**
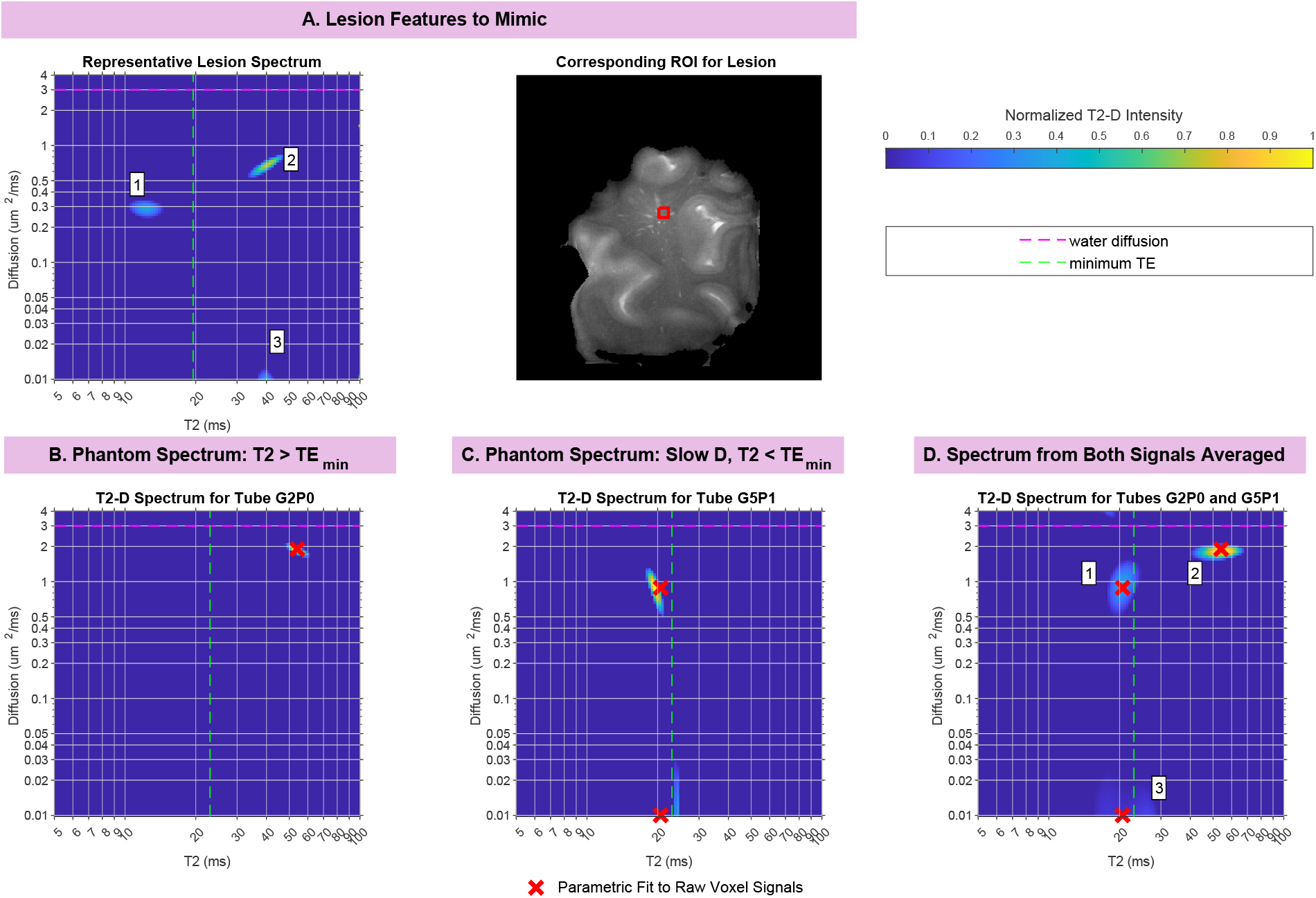
Phantom application to replicate spectral components observed in lesioned tissue. (A) Representative fast spin echo image with the analyzed lesion ROI highlighted (right) and corresponding T_2_–D spectrum (left). (B) Single-component T_2_–D spectrum from tube G2P0, characterized by a T_2_ exceeding the TE_min_. The signal’s parametric fit is marked by 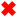. (C) Two-component T_2_–D spectrum from tube G5P1, exhibiting similar T_2_ values but diffusion coefficients differing by approximately two orders of magnitude. The two 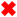 marks represent the bi-exponential parametric fit for the individual tube’s signal. (D) Signals (not spectra) from tubes G2P0 and G5P1 were combined and processed with NNLS to generate a composite spectrum with component T_2_ and D values comparable to those observed in lesion ROIs. These results demonstrate that phantom configurations incorporating short-T_2_ and restricted-diffusion components reproduce key spectral characteristics observed in lesioned tissue. The three 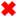 marks are translated from B and C for reference.

In the composite, phantom-derived spectrum from tubes G5P1 and G2P0, all components retained their relative positions. The slow diffusion or short T_2_ features did not prevent the resolution of coexisting components under the acquisition and reconstruction conditions.

#### NAWM spectrum validation

We were able to use phantom tube G7P1 (Figure 4B) to derive a component with a short-T_2_, which was less than half the TE_min_, similar to the NAWM’s Component 1 (Figure 4A). As this tube also contained PEG, it exhibited a faint secondary component at a similar T_2_, but with substantially slower diffusion. Components 2 and 3 from the NAWM were replicated using tube G3P1 (Figure 4C).

**FIGURE 4.**
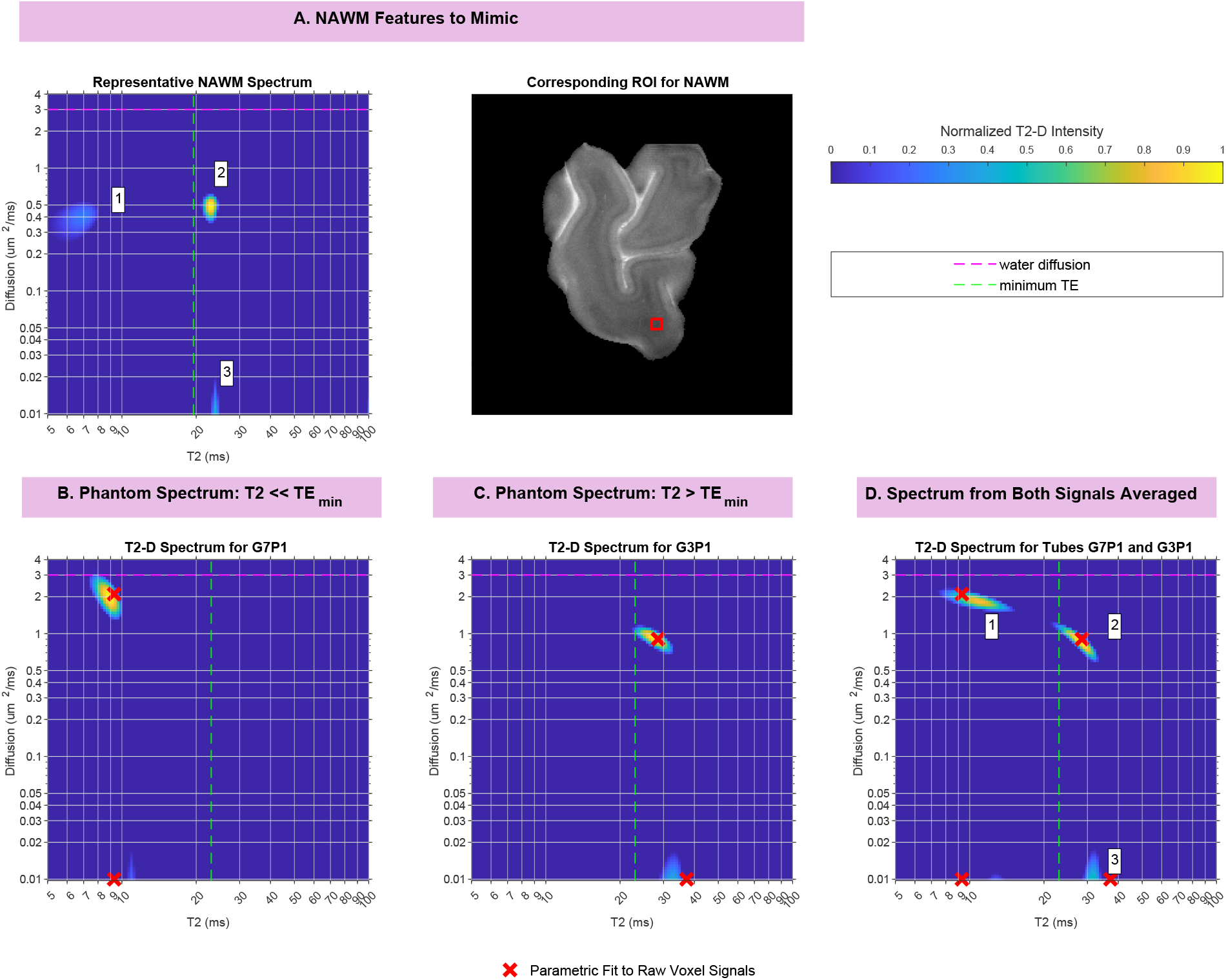
(A) Representative fast spin echo image with the analyzed NAWM ROI highlighted (right) and corresponding T_2_–D spectrum (left). (B) Two-component T_2_–D spectrum from tube G7P1 containing two components with T_2_ less than half the TE_min_. The two 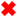 marks represent the bi-exponential parametric fit for the tube’s signal. (C) Two-component T_2_–D spectrum from tube G3P1 containing two components with T_2_ above the TE_min_. (D) The composite spectrum shows three peaks with similar relative positions with respect to the TEmin and the lowest resolvable diffusion coefficient. These results demonstrate the feasibility of detecting components with T_2_s expected for healthy myelin. The four 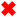 marks are translated from B and C for reference.

The combined spectrum resolved three distinct components with centroids consistent with those observed in the corresponding single-tube spectra (Figure 4D). Therefore, allowing our NNLS algorithm to fit the NAWM component 1, speculated to be myelin, did not significantly impact the other components.

### 3.3 Simulations

Figure 5 summarizes the effects of TE_min_ and SNR on the diffusion and T_2_ coefficients resolved from the simulated CDR data. Increasing TE_min_ and decreasing SNR resulted in progressive elongation of the components along the diffusion dimension, as quantified by the diffusion peak spread (Figure 5A). Substantial broadening 1-2 orders of magnitude relative to an ideal discrete component was observed primarily at the lowest tested SNR (10 dB) and for TE_min_ values exceeding 25 ms.

**FIGURE 5.**
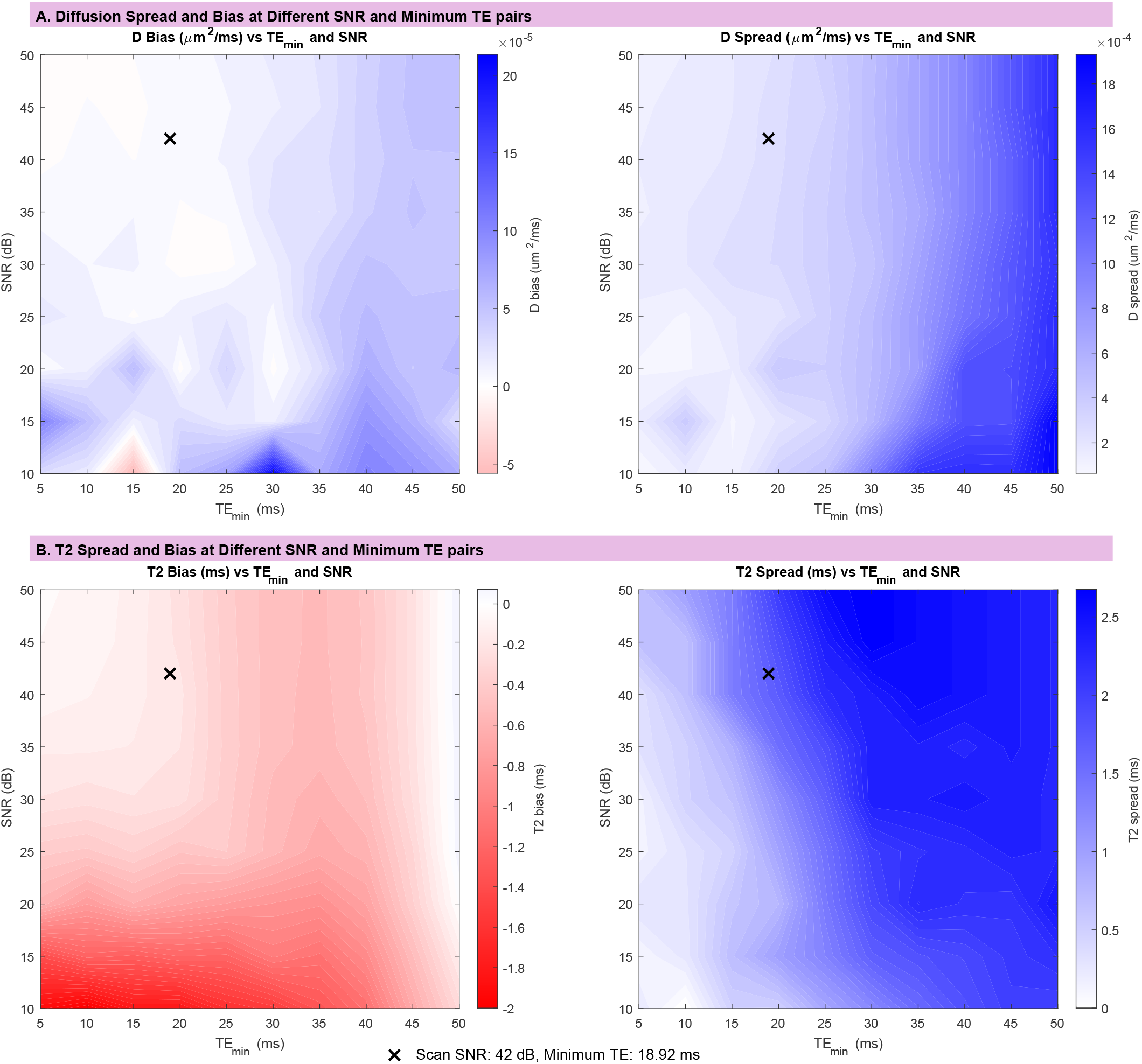
Simulations evaluating effect of SNR and TE_min_ on T_2_–D spectral accuracy. The black *×* denotes the SNR (42 dB) and TE_min_ (18.92 ms) used in the tissue scanning protocols. A) Diffusion bias and spread as a function of SNR and TE_min_: Contour plot (left) showing displacement of the resolved diffusion centroid relative to the ground-truth diffusion coefficient (bias), with red indicating negative bias and blue indicating positive bias. Contour plot (right) illustrating elongation (effective diameter or spread) of the resolved spectral peak along the diffusion axis as a function of SNR and TE_min_. B) T2 bias and spread as a function of SNR and TE_min_: Contour plot (left) depicting the resolved T_2_ centroid bias from the ground-truth relaxation time as SNR and TE_min_ vary. Contour plot (right) demonstrating the effects of SNR and TE_min_ on the spread of the resolved spectral peak along the T_2_ axis. For the experimentally relevant TE_min_ and measured SNR (black *×*), both diffusion and T_2_ centroids remain minimally biased.

The diffusion bias plot (Figure 5B) indicates that, across conditions, the diffusion centroid appeared either faster or slower than the true diffusion value. At higher SNRs combined with longer TE_min_, the diffusion centroid was biased toward faster apparent diffusion coefficients. Conversely, for short TE_min_ values (5–10 ms) and reduced SNR, the centroid shifted toward more restricted diffusion relative to the ground truth. As TE_min_ increased beyond this ideal range, the apparent diffusion progressively overestimated the true diffusion coefficient.

Along the T_2_ dimension, the centroid systematically shifted toward shorter T_2_ values with increasing TE_min_ and decreasing SNR (Figure 5C,D). Reductions in SNR produced a more pronounced effect on T_2_ centroid bias than increases in TE_min_. For SNR values below 20 dB, the T_2_ centroid deviated by approximately 1–2 ms toward shorter relaxation times.

## 4 DISCUSSION

We evaluated the feasibility and accuracy of using CDR at 7T for studying WM spectral features in ex vivo ADRD tissue. We first briefly characterized NAWM and lesioned tissue T_2_–D spectral components, recreated them with phantoms to identify unwanted component interactions and artifacts, and ran simulations to determine if the practical achievable SNR and TE_min_ can capture WM features of interest.

Our ex vivo tissue CDR analysis consistently exhibited multiple components, including short-T_2_ and restricted-diffusion features. The results align with prior human WM T_2_–D studies at 3T ^30^ and 7T ^21,33^ that also report consistent multi-component spectra. Differences in T_2_ and diffusion coefficients across studies are plausibly explained by physiology (in vivo vs ex vivo) and tissue preparation (fixed vs thawed).

Our phantom allowed us to emulate T_2_–D components that were observed in NAWM and lesioned tissue. Through the iterative process of creating the composite spectra, we found that the component speculated to be myelin water does not significantly bias the estimations of the axonal or EC water features.

Finally, based on simulations, we found that reliable short-T_2_ recovery required SNR *>* 30 dB and TE_min_ *<* 25 − 30 ms. When SNR is lower or TE_min_ is longer, centroid bias and peak broadening increase, and lowamplitude features near the TE_min_ limit warrant greater scrutiny.

Limitations include our small tissue sample size and use of a single diffusion encoding direction. Anisotropic tissue will require multiple directions (e.g., for powder-averaging), ^24^ which will increase scan time. Future studies include optimizing the number and placement of TE and *b* value combinations, extending to a larger cohort, and comparing spectra results with histology of the tissue.

For future studies, we suggest: **1)** evaluating SNR to ensure the fitting routine is not resolving components biased by noise, **2)** using the component centroid T_2_–D values to interpret the spectra, not the spread, and **3)** treating features near the TE_min_ boundary and low-intensity components as candidates requiring additional validation.

## 5 CONCLUSION

This work used phantom and simulation-based validation to evaluate short-T_2_ and restricted-diffusion components observed in combined T_2_–diffusion MRI of ex vivo ADRD WM. Across NAWM, lesions, and GM, three consistent spectral components were observed, including a component with T_2_ shorter than the TE_min_ and a component with highly restricted diffusion.

Phantom experiments demonstrated that components with T_2_ values around 10 ms and components near the lower bound of diffusion sensitivity were recoverable and did not induce spurious spectral components when combined with longer-T_2_ compartments. Complementary simulations showed that, under experimentally relevant SNRs and TE_min_s, spectral centroids remained minimally biased and peak broadening remained limited.

Together, these results indicate that the observed short-T_2_ and restricted-diffusion components are unlikely to arise solely from acquisition limitations. Instead, they reflect resolvable and reproducible features of the underlying signal, supporting their interpretation as biologically meaningful signatures that could be consistent with myelin-associated and axonal water compartments in ADRD WM.

## ACKNOWLEDGMENTS

We thank Ulrich Scheven for his contributions to helpful discussions on CDR and guidance scanning on the 7T Varian animal scanner.

